# Cardioprotection by Poloxamer 188 is Mediated through Increased Endothelial Nitric Oxide Production

**DOI:** 10.1101/2024.05.18.593838

**Authors:** Gaoxian Chen, Hunter F. Douglas, Zhu Li, William J. Cleveland, Claudius Balzer, Demetris Yannopolous, Ian Ying-Li Chen, Detlef Obal, Matthias L. Riess

## Abstract

Ischemia/reperfusion (I/R) injury significantly contributes to the morbidity and mortality associated with cardiac events. Poloxamer 188 (P188), a nonionic triblock copolymer, has been proposed to mitigate I/R injury by stabilizing cell membranes. However, the underlying mechanisms remain incompletely understood, particularly concerning endothelial cell function and nitric oxide (NO) production.

We employed human induced pluripotent stem cell (iPSC)-derived cardiomyocytes (CMs) and endothelial cells (ECs) to elucidate the effects of P188 on cellular survival, function, and NO secretion under simulated I/R conditions. iPSC-CMs contractility and iPSC-ECs’ NO production were assessed following exposure to P188. Further, an isolated heart model using Brown Norway rats subjected to I/R injury was utilized to evaluate the ex-vivo cardioprotective effects of P188, examining cardiac function and NO production, with and without the administration of a NO inhibitor.

In iPSC-derived models, P188 significantly preserved CM contractile function and enhanced cell viability after hypoxia/reoxygenation. Remarkably, P188 treatment led to a pronounced increase in NO secretion in iPSC-ECs, a novel finding demonstrating endothelial protective effects beyond membrane stabilization. In the rat isolated heart model, administration of P188 during reperfusion notably improved cardiac function and reduced I/R injury markers. This cardioprotective effect was abrogated by NO inhibition, underscoring the pivotal role of NO. Additionally, a dose-dependent increase in NO production was observed in non-ischemic rat hearts treated with P188, further establishing the critical function of NO in P188 induced cardioprotection.

In conclusion, our comprehensive study unveils a novel role of NO in mediating the protective effects of P188 against I/R injury. This mechanism is evident in both cellular models and intact rat hearts, highlighting the potential of P188 as a therapeutic agent against I/R injury. Our findings pave the way for further investigation into P188’s therapeutic mechanisms and its potential application in clinical settings to mitigate I/R-related cardiac dysfunction.

## Introduction

More than 380,000 patients in the United States are victims of out-of-hospital cardiac arrest each year [42]. Even with the most effective cardiopulmonary resuscitation (CPR), more than 90 percent of patients suffer serious neurological damage or die [9]. Since only a minority receives bystander CPR, and first responders arrive on average 8-10 min later, abrupt reintroduction of blood flow at the initiation of CPR after prolonged ischemia has additional detrimental effects on tissue recovery and potentiates the resulting ischemia/reperfusion (I/R) injury [34].

The intricate interplay between cellular membranes and pathological stress responses, particularly during I/R injury, has garnered considerable attention in the realm of biomedical research. Copolymer-based cell membrane stabilizers (CCMS), comprising hydrophobic poly-propylene oxide (PPO) and hydrophilic poly-ethylene oxide (PEO) chains, have emerged as potent agents in mitigating membrane damage and curtailing cell death pathways [3, 35, 36, 66]. Among these, Poloxamer 188 (P188, E_75_P_30_E_75_, Sigma Aldrich), a non-ionic tri-block copolymer, has been spotlighted for its ability to bridge gaps in compromised cell membranes [37], thereby obstructing the formation of detrimental membrane pores under conditions of intense cellular stress experienced during ischemia [36]. The protective mechanism of P188 extends beyond mere physical membrane repair; it encompasses the prevention of unregulated ion flux across cellular boundaries and the blockade of apoptosis by maintaining calcium equilibrium and preventing mitochondrial depolarization under oxidative stress [58, 66]. This dual action not only safeguards skeletal muscle and neurons from I/R-induced damage but also fortifies the blood-brain barrier against various cerebral challenges, significantly reducing brain edema and neuronal cell death [1, 18, 32, 40, 52, 60, 61]. Both, *in-vivo* and *ex-vivo* models of cardiac I/R injury demonstrated the protective effect of P188 against cardiomyocyte (CM) cell necrosis and apoptosis when given immediately upon the time of reoxygenation [35].

Building on the foundational role of the vascular endothelium in I/R injury, our research delves into the nuanced mechanisms by which endothelial cells (ECs) orchestrate a protective response against such injuries, specifically through the modulation of nitric oxide (NO) production [8, 14, 38, 51]. NO stabilizes calcium homeostasis and attenuates calcium overload in CMs [7, 39], activating intracellular protein kinase pathways that inhibit mitochondrial permeability transition pore opening [13, 33, 62], scavenging reactive oxygen species and suppressing oxidative damage, optimizing oxygen metabolism [43] and energy production in CMs [56, 65]. This complex cascade of protective measures is initiated by flow-induced shear stress, which activates endothelial mechanosensors such as VEGFR2, PECAM-1, and integrins [6, 44, 49, 54, 59], leading to an increase in NO production through phosphorylation [11, 15] and increased expression of endothelial NO synthase (eNOS) [10]. To empirically test our hypothesis, we employed a multifaceted experimental design encompassing human induced pluripotent stem cell-derived CMs (iPSC-CMs) and iPSC-ECs, as well as rat isolated, intact hearts. Our findings indicate that P188 not only enhances the contractile function of iPSC-CMs but also significantly increases the viability of both iPSC-CMs and iPSC-ECs following hypoxia/reoxygenation. A notable aspect of our study is the observed increase in NO secretion specifically in iPSC-ECs, a finding that underscores the role of P188 in modulating EC function in a manner that is beneficial for cardiac recovery after I/R injury. This endothelial-centric protective effect was further corroborated in rat isolated hearts, where P188 administration was associated with improved cardiac function and viability after ischemia, in a manner dependent on NO signaling. Intriguingly, P188 was also found to elevate NO levels in non-ischemic isolated hearts in a dose-dependent manner, suggesting a broader role for P188 in cardiovascular protection that extends beyond its immediate effect on membrane integrity.

Collectively, our research presents a novel paradigm in the understanding of cardiac protection against I/R injury, positioning NO not merely as a bystander but as a central player in the protective effects mediated by P188. Through a detailed exploration of the interaction between P188, ECs, and CMs, we provide compelling evidence for a synergistic mechanism of action that holds considerable promise for the development of novel therapeutic strategies against myocardial I/R injury.

## Materials and Methods

### Human induced pluripotent stem cells (iPSC)

The iPSC line was obtained from the Stanford Cardiovascular Institute Biobank (http://med.stanford.edu/scvibiobank.html). Patients were enrolled under the Stanford Institutional Review Board and Stem Cell Research Oversight Committee guidelines. Patients consent was retrieved and the iPSCs were generated from peripheral blood mononuclear cells (PBMCs) using Sendai virus (CytoTuneTM-iPS 2.0 Sendai Reprogramming Kit, Thermo Fisher Scientific) as previously described. Differentiation of iPSCs into iPSC-ECs [45, 63] and -CMs [5] followed previously published protocols (***Fig. 1A top panel, Fig. 3A***).

**Fig. 1.**
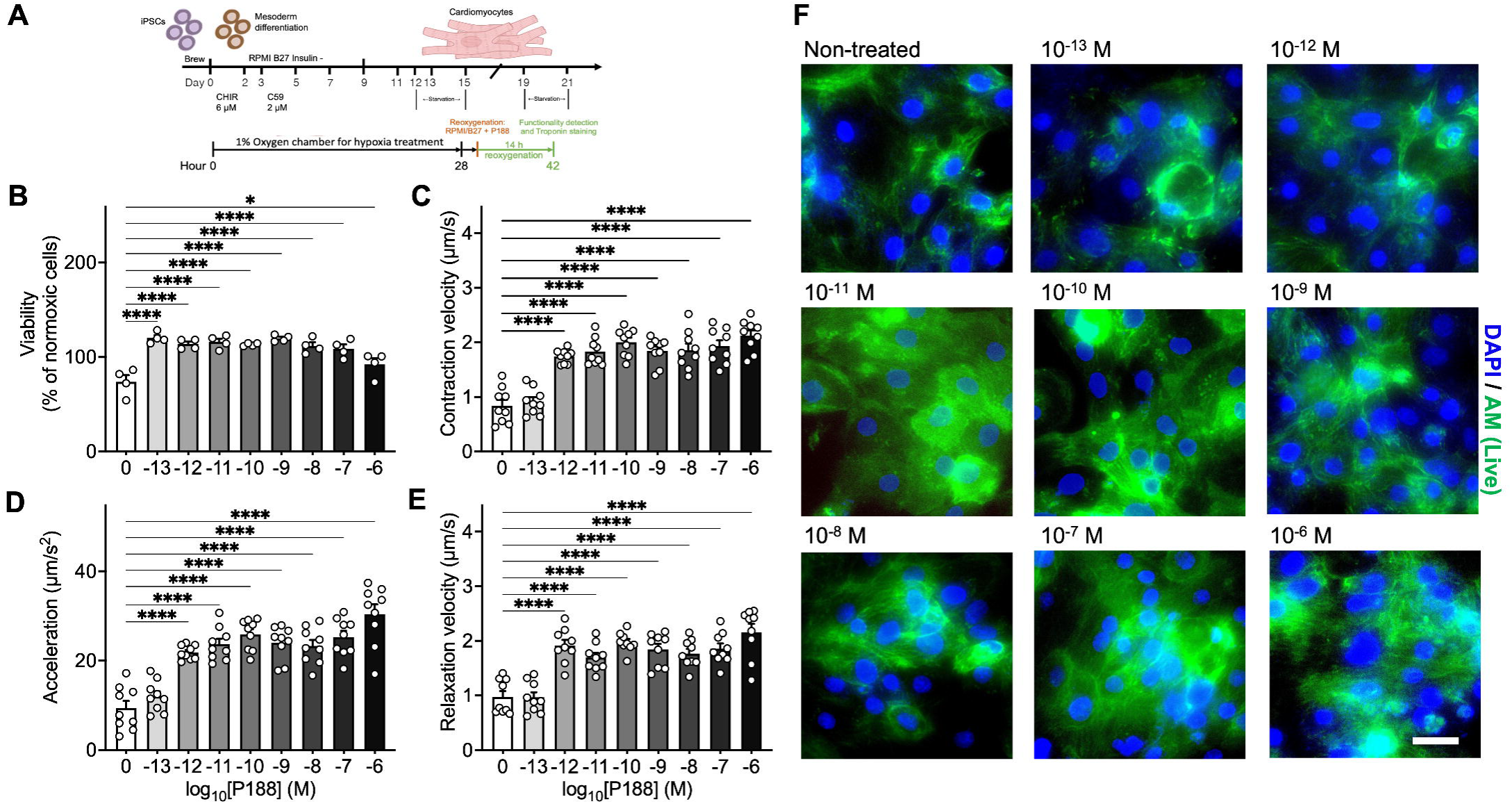
P188-mediated protection of iPSC-CMs under simulated H/R conditions. (A) Schematic diagram illustrating the differentiation process of iPSCs to iPSC-CMs, followed by hypoxic challenge and subsequent reoxygenation phase supplemented with various P188 concentrations. (B) Quantitative assessment of iPSC-CM viability across different P188 concentrations under simulated H/R conditions. The data emphasizes a marked increase in cell survival with increasing P188 concentrations. Cell viability values are normalized to normoxic control cells. (C-E) Quantitative representation showcasing the augmented functionalities of iPSC-CMs, including contraction rate (C), relaxation rate (D), and acceleration (E), upon P188 exposure under simulated H/R conditions. (F) Representative images depicting the distribution of troponin T in iPSC-CMs treated with varying P188 concentrations. Staining was shown as nuclei in blue and troponin in green; scale bar, 30 μm. Data are shown as mean ± SEM. Significance levels in comparison to the non-treated control group were determined using one-way ANOVA with Dunnett’s test: *P<0.05, ****P<0.0001.

### Differentiation of iPSC-CMs

iPSCs were next cultured to 85% cell confluence, and then treated for 2 days with the GSK 3 inhibitor CHIR99021 (6 μM, Selleck Chemical) in RPMI medium plus B27 supplement without insulin medium to activate WNT signaling and induce mesodermal differentiation. On day 2, cells were placed on RPMI medium plus B27 supplement without insulin medium and CHIR99021 (3 μM) [5, 27]. On days 3-4, cells were treated with C59 (2 μM, Selleck Chemical) to inhibit the WNT pathway and induce cardiogenesis. On days 5-6, C59 was removed, and cells were cultured in RPMI medium plus B27 supplement without insulin medium. From day 7 onwards, cells were cultured in RPMI medium plus B27 supplement with insulin medium until cardiac contractions were observed. At this point, cells were cultured for 4 days in glucose-free RPMI medium plus B27 supplement (Gibco) with insulin medium to purify iPSC-CMs. Following purification, cells were cultured in RPMI medium plus B27 with insulin medium which regular media changes every other day (***Fig. 1A, top panel***). When re-plating iPSC-CMs for downstream use, cells were dissociated with Trypsin/EDTA (0.25%, Life Technologies) into a single-cell suspension and seeded on Matrigel-coated plates with ROCK inhibitor and knockout serum replacement (1%, Gibco).

### Differentiation of iPSC-ECs

Briefly, iPSCs were dissociated using 0.5 mM EDTA/ phosphate-buffered saline (PBS, Innovative Cell Technologies) and plated as single cells in Stem cell brew (Miltenyi Biotec) supplemented with ROCK inhibitor Y-27632 (Selleck Chemical) on Matrigel to a final density of 20,000-30,000 cells/cm^2^. iPSC monolayers were cultured to 85% cell confluency. iPSCs were then treated with the glycogen synthase kinase (GSK) 3 inhibitor CHIR99021 (6 μM, Selleck Chemical) for 2 days in RPMI medium plus B27 supplement without insulin (Gibco, CA). On day 2, cells were treated for 48 hours with a lower dose of CHIR99021 (2 μM) in RPMI medium plus B27 supplement without insulin medium (Gibco). On day 4, cells were treated with the recombinant human fibroblast growth factor-2 (rhFGF-2, 10 ng/ml, PeproTech) and the vascular endothelial growth factor (VEGF, 20 ng/ml, R&D Systems) for EC expansion for 8 days. On day 12, cells were harvested with Trypsin/EDTA (0.25%, Life Technologies) and sorted using anti-CD144 MicroBeads (Thermo Fisher Scientific). The differentiation efficiency was calculated by the number of CD144-positive cells (***Fig. 3A***). iPSC-ECs were cultured in gelatin-coated (Sigma Aldrich) 6-well plates in EGM-2 medium (Lonza).

### Hypoxia/reoxygenation experiments in iPSC-CMs and iPSC-ECs

iPSC-CMs were seeded at 30,000 cells per well on 384-well plates and cultured for 96 hours prior to hypoxic/reoxygenation (H/R) treatment. iPSC-ECs were seeded on 24-well plates and cultured until reaching 95% confluence (2-3 days). To induce significant cell death (∼50%), iPSC-CMs were exposed to 28 hours of hypoxia (FiO_2_ = 0.01, measured with an FD-90A-O_2_ oxygen meter) followed by 14 hours of reoxygenation with varying P188 concentrations (***Fig. 1A, lower panel***). iPSC-ECs were exposed to 36 hours of hypoxia followed by 5-8 hours of reoxygenation (***Fig. 3B***). The effects of P188 treatment on iPSC-EC viability, NO production, and CM functionalities were subsequently characterized.

### Cell viability assays in iPSC-CMs and iPSC-ECs

Following hypoxic treatment, iPSC-CMs and iPSC-ECs were reoxygenated in RPMI plus supplement and insulin medium containing different concentrations of P188 for 14 hours and 5 hours, respectively. After reoxygenation with P188, cell viability was assessed by adding an equal volume of Titer-Glo reagent (Promega) to the medium and incubating for 3 minutes at room temperature. Plates were preheated to 37°C for 7 minutes before measuring luminescence on a Promega GloMax Multi plate reader. Relative luminescence units (RLU) were recorded to quantify cell viability.

### Apoptosis of iPSC-CM detection by flow cytometry

Apoptosis was detected using the Annexin V-FITC/PI Apoptosis Detection Kit (Vazyme, A211) according to the manufacturer’s instructions. Briefly, cells (5 × 10^5^) were harvested, washed twice with PBS, and resuspended in 1X Binding Buffer. The cell suspension was incubated with 5 μl Annexin V-FITC and 5 μl PI Staining Solution for 10 minutes at room temperature in the dark. After incubation, 400 μl of 1X Binding Buffer was added, and the stained cells were analyzed within one hour on a table-top flow cytometry (BD C6 Accuri Flow Cytometer) using the 488 nm laser for excitation and the FL1 channel (FITC) and FL3 channel (Propidium Iodide [PI], Invitrogen) for detection. A total of 10,000 events were collected for each sample. Data analysis was performed using FlowJo software (version 10.8.1). The cells were classified into three subpopulations based on Annexin V-FITC and PI staining: viable cells (AnnexinV-FITC^-^/PI^-^), early apoptotic cells (AnnexinV-FITC^+^/PI^-^), and late apoptotic/necrotic cells (AnnexinV-FITC^+^/PI^+^).

### Measurement of iPSC-CM contractility

The contractility of iPSC-CMs was assessed after 14 hours of reoxygenation using a cell motion imaging system (Sony SI8000, Sony Biotechnology). Videos of beating iPSC-CMs were recorded for 10 seconds at 75 frames per second (fps) with a resolution of 2048×2048 pixels using a 10× objective lens. CM contraction was analyzed using Sony SI8000 analysis software as previously described [55].

### Measurement of nitric oxide production in iPSC-ECs

Intracellular NO production was assessed during reoxygenation by live-cell staining with the NO-sensitive fluorescent dye 4-Amino-5-Methylamino-2’,7’-Difluorofluorescein (DAF-FM) diacetate (Invitrogen). iPSC-ECs were stained with 1 μM DAF-FM diacetate for 10 minutes at 37°C, in the last 10 minutes of reoxygenation at 7 hours 50 minutes after reoxygenation commenced. Subsequently, Hoechst 33342 (1 μg/mL, Thermo Scientific) was added to label nuclei. Imaging was performed on a Revolve microscope (Echo Laboratories) under 10× magnification. DAF-FM signals were detected in the GFP channel and Hoechst signals in the DAPI channel. DAF-FM fluorescence was also quantified using the GFP channel of a Cytation5 imaging system (BioTek, Agilent Technologies).

### Rat isolated heart experiments

#### Animals

All investigations abided by the Guide for the Care and Use of Laboratory Animals (Institute for Laboratory Animal Research, National Academy of Sciences, 8th edition, 2011) and were previously approved by the Institutional Animal Care and Use Committees of Vanderbilt University Medical Center and the Tennessee Valley Healthcare System - Veterans Affairs Medical Center (both in Nashville, TN, USA). Juvenile (8-10 weeks old) male Brown Norway (BN) rats were obtained from Charles River Laboratories International, Inc. The BN rat was chosen as it has been identified to be the rat strain most resistant against myocardial I/R injury, and produces higher level of endogenous NO [2, 23]. A total of 40 BN rats were used to complete the studies.

#### Langendorff heart preparation

Animals were anesthetized with intraperitoneal injection of 100 mg/kg of 100 mg/mL ketamine (Hospira) followed by intraperitoneal injection of 3,000 U/kg heparin (Fresenius Kabi). After a negative response to a noxious stimulus 10 minutes later, animals were decapitated and hearts were excised via thoracotomy, following rapid cannulation of the aorta, distal to the aortic valve, and ligation of the inferior and superior venae cavae. The hearts were perfused retrograde with 4 °C cold oxygenated Krebs-Henseleit solution with the following composition (in mM): 148 Na^+^, 4.7 K^+^, 1.2 Mg^2+^, 1.6 Ca^2+^, 127 Cl^−^, 27.8 HCO_3_^−^, 1.2 H_2_PO_4_^−^, 1.2 SO_4_^2−^, 5.5 glucose, 2 pyruvate, 0.026 ethylene diamine tetraacetic acid, and 5 U/L insulin. The heart was promptly placed into the support system and perfused at 70 mmHg and 37 °C. The perfusate was equilibrated with 95% O_2_ and 5% CO_2_ to maintain a constant pH of 7.40 (carbon dioxide partial pressure pCO_2_ 40 mmHg; oxygen partial pressure pO_2_ ∼570 mmHg). The perfusate was filtered using 5 µm pore size filters in-line. Left ventricular pressure (LVP) was measured isovolumetrically via a saline-filled latex balloon (Radnoti LLC, Monrovia, CA) inserted directly into the left ventricle by way of the left atrium. The initial volume of the balloon was titrated to achieve a diastolic LVP of 10 mmHg at baseline so that any consequent increase reflected diastolic contracture. LVP-derived data included systolic (LVSP), diastolic (LVEDP), and developed (systolic minus diastolic) LVP (LVDP), and its maximum and minimum first derivatives (dP/dt_max_ and dP/dt_min_) as indicators of ventricular contractility and relaxation, respectively. Spontaneous heart rate (HR) was monitored via electrocardiogram using bipolar electrodes placed in the right atrial and right ventricular walls. The rate pressure product (RPP) was calculated as developed LVP × HR to correct for HR-induced decreases in LVDP due to changes in sarcoplasmic reticulum calcium release. Changes in coronary flow were measured by an in-line ultrasonic flowmeter (T106X; Transonic Systems).

#### Ischemia/reperfusion studies in rat isolated hearts

Langendorff prepared hearts were randomized to reperfusion with vehicle (control, n=5), 1.0 mM P188 (n=5), 100 µM of the nonspecific nitric oxide synthase inhibitor Nω-Nitro-L-arginine methyl ester hydrochloride (L-NAME [Sigma Aldrich], n=5), or 1.0 mM P188 and 100 µM L-NAME (n=5). 1.0 mM of the polymer was chosen following a series of preliminary I/R studies using 0.1 mM P188, 0.3 mM P188, 1.0 mM P188, or 3.0 mM P188. Additionally, our prior experiments in the *ex-vivo* model of myocardial I/R have identified 1 mM as the optimal concentration to elicit cardioprotection. Hearts were allowed to stabilize for 25 minutes. Following baseline readings, hearts were subjected to 30 minutes of global no-flow ischemia, immediately followed by 120 minutes of reperfusion with continuous monitoring of LVP, heart rate, and coronary flow, followed by tissue harvest and ventricular infarct size assessment. If ventricular fibrillation occurred during reperfusion, a bolus of lidocaine (250 µg) was immediately injected into the aortic cannula. All data were collected from hearts naturally in, or converted to, sinus rhythm.

#### Infarct size measurement

At the end of each I/R experiment, hearts were removed from the system, and weighed after the removal of both atria. Using a heart matrix, the ventricles were cut into 2-mm transverse sections. Heart slices were then incubated with 2,3,5-triphenyltetrazolium chloride (1%, TTC, Sigma Aldrich) in KH_2_PO_4_ buffer (100 mM, pH 7.4, 37°C) for 10 min to stain viable tissue red by dehydrogenase enzymes present in viable cells, with infarcted areas remaining white [48]. Each heart slice was digitally imaged on a green background, and their infarcted areas were measured via planimetry using Image J (Ver 2.14.0, NIH, Bethesda, MD) software, its ColorThreshold plugin, and a custom-developed macro to ensure unbiased measurements[53]. Infarcted areas were averaged based on their weight to calculate the total ventricular infarction of each heart [48].

#### Nitric oxide production studies in normoxic rat isolated hearts

Non-ischemic Langendorff-prepared hearts were randomized to perfusion with either vehicle only (control, n=3), or P188 (n=3). Hearts were allowed to stabilize for 25 minutes, then loaded with 10 μM of the NO-specific DAF-FM. P188 was administered in increasing concentrations (0 mM, 0.1 mM, 0.3 mM, 1.0 mM, and 3.0 mM) for 10 minutes; vehicle control experiments were conducted for the same duration but without P188 infusion. DAF-FM fluorescence was subsequently measured using a spectrophotometer via a bifurcated fiber optic probe in contact with the left ventricular epicardial wall. Fluorescence was excited at 495 nm and emitted at 515 nm [46]. This experimental procedure was repeated in the same hearts after the addition of 100 μM L-NAME to each concentration of P188 and control. Thereafter, 10 µM of the exogenous NO donor sodium nitroprusside was added to perfusate to ensure the fluorescent signal was not saturated (not shown). Since DAF fluorescence is irreversible, i.e. additive, fluorescence increases from before, not the absolute values, were used as a surrogate of NO release.

#### Statistical analysis

All analog signals recorded throughout the studies were digitized (NI USB-6343, National Instruments Corporation) and recorded at 200 Hz (Labview, National Instruments) for later analysis. All data were tested for normal distribution and equal variance. If the data did not pass both tests, we analyzed them by non-parametric testing with either the Kruskal-Wallis or the Mann-Whitney test as appropriate. If they did pass both tests, data were compared by parametric one-way analysis of variance. The Student-Newman-Keul test was used for post-hoc testing among the groups. Since not all data were normally distributed and had equal variance, we display all of them as box plots with median and interquartile range for consistency, unless otherwise indicated. For cell-based assays we determined differences compared to the control group using Dunnet’s post hoc test; they are displayed as means ± standard error. Tests were considered statistically significant at P<0.05 (one symbol; P<0.01 two symbols; P<0.001 three symbols; P<0.0001 four symbols): *vs CON/control, † vs ISC, ‡ vs P188, § vs P188 + L-NAME.

## Results

### iPSC-derived cell studies

#### iPSC-CM function after hypoxia/reoxygenation experiments

iPSC-CMs were subjected to hypoxia by exposing them to 1% O_2_ for 28 hours in a glucose-deprived medium to mimic hypoxic conditions. Following the hypoxia phase, iPSC-CMs were provided with a glucose-containing and oxygen-rich medium supplemented with varying concentrations of P188 to assess its protective effects (***Fig. 1A***). Remarkably, the protective effects of P188 on iPSC-CMs increased with higher concentrations, achieving nearly 90% cell survival (***Fig. 1B***), in contrast to the 50% survival in the untreated group. Moreover, P188 treatment significantly enhanced iPSC-CM function, including contraction rate, relaxation rate, and acceleration (***Fig. 1C-E***), suggesting its role in preserving and recovering iPSC-CM function under hypoxic conditions. The morphological evaluation of iPSC-CMs at different P188 concentrations (***Fig. 1F***) revealed that untreated cells exhibited cell shrinkage and an unhealthy morphology, while P188 at concentrations ranging from 1 µM to 100 fM effectively maintained healthy and intact cellular morphology.

#### iPSC-derived CM cell viability after hypoxia/reoxygenation experiments

P188 restored intracellular adenosine triphosphate (ATP) levels after 14 hours of reoxygenation as measured by the Cell titer glue assay (***Fig. 1B***). This effect was independent of the dose of P188 and might reflect the restored cellular metabolism in cells exposed to P188. In contrast, the cell live-dead staining indicated a dose-dependent protective effect of P188 against reoxygenation-induced injury in iPSC-CMs (***Fig. 2 A, 2B***).

**Fig. 2.**
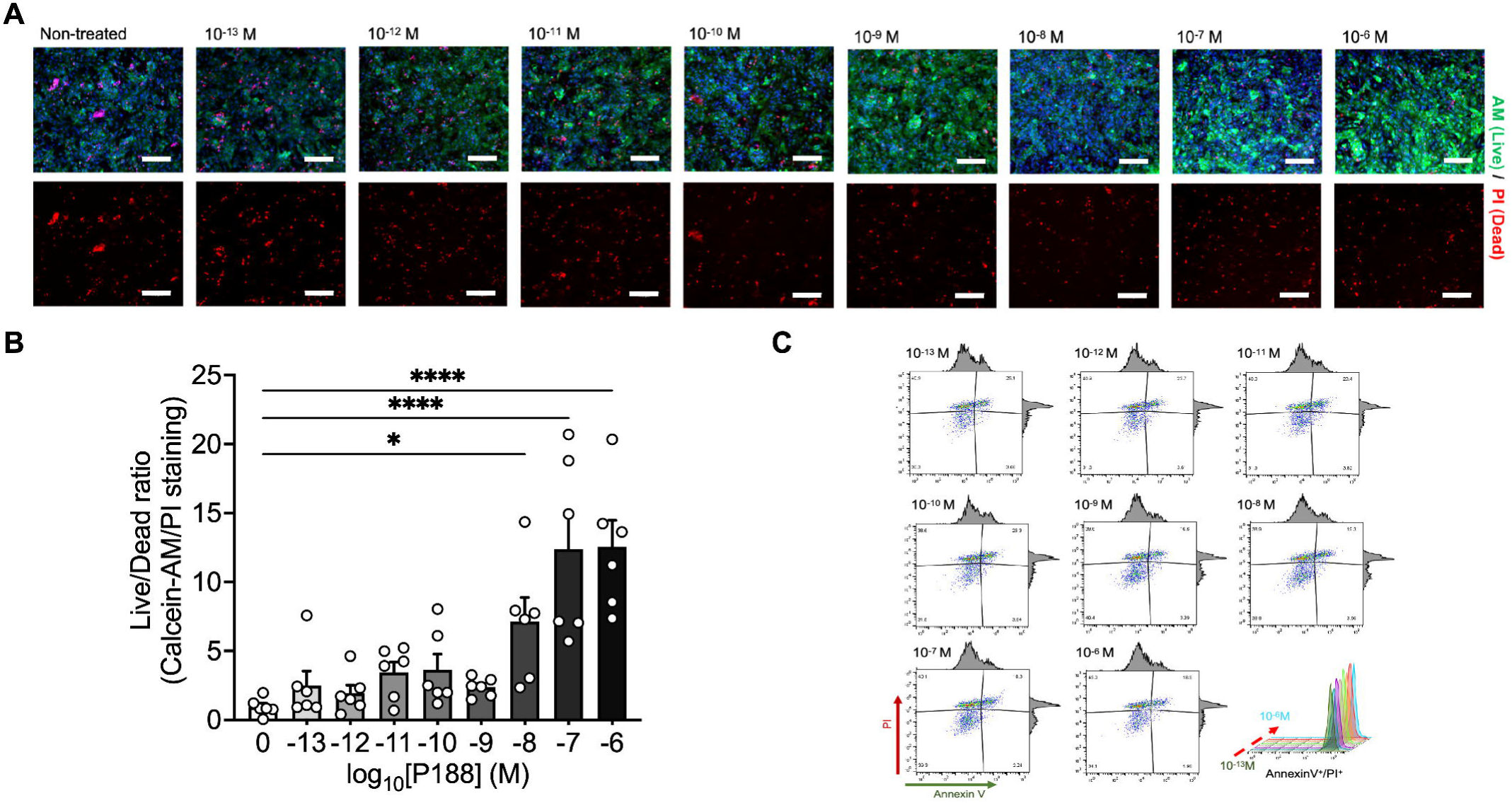
Cell death analysis and apoptosis detection in iPSC-CM. (A) Live/dead staining with Calcein-AM (green) and Propidium Iodide (PI, red) of iPSC-CMs after treatment with a concentration series of P188 ranging from 10^-13^ M to 10^-6^ M. Non-treated samples serve as controls. Scale bars, 130 μm. (B) Quantification of the live/dead cell ratio in iPSC-CMs following reoxygenation, with or without P188 treatment. (C) Flow cytometry analysis of Annexin V and PI staining for detection of apoptosis in iPSC-CMs. The stacked peaks represent the population of Annexin V-positive/PI-positive cells, indicating an increasing ratio of late apoptotic/necrotic cells in P188-treated iPSC-CMs. Data are shown as mean ± SEM. Significance levels in comparison to the non-treated control group were determined using one-way ANOVA with Dunnett’s test: *P<0.05, ****P<0.0001.

**Fig. 3.**
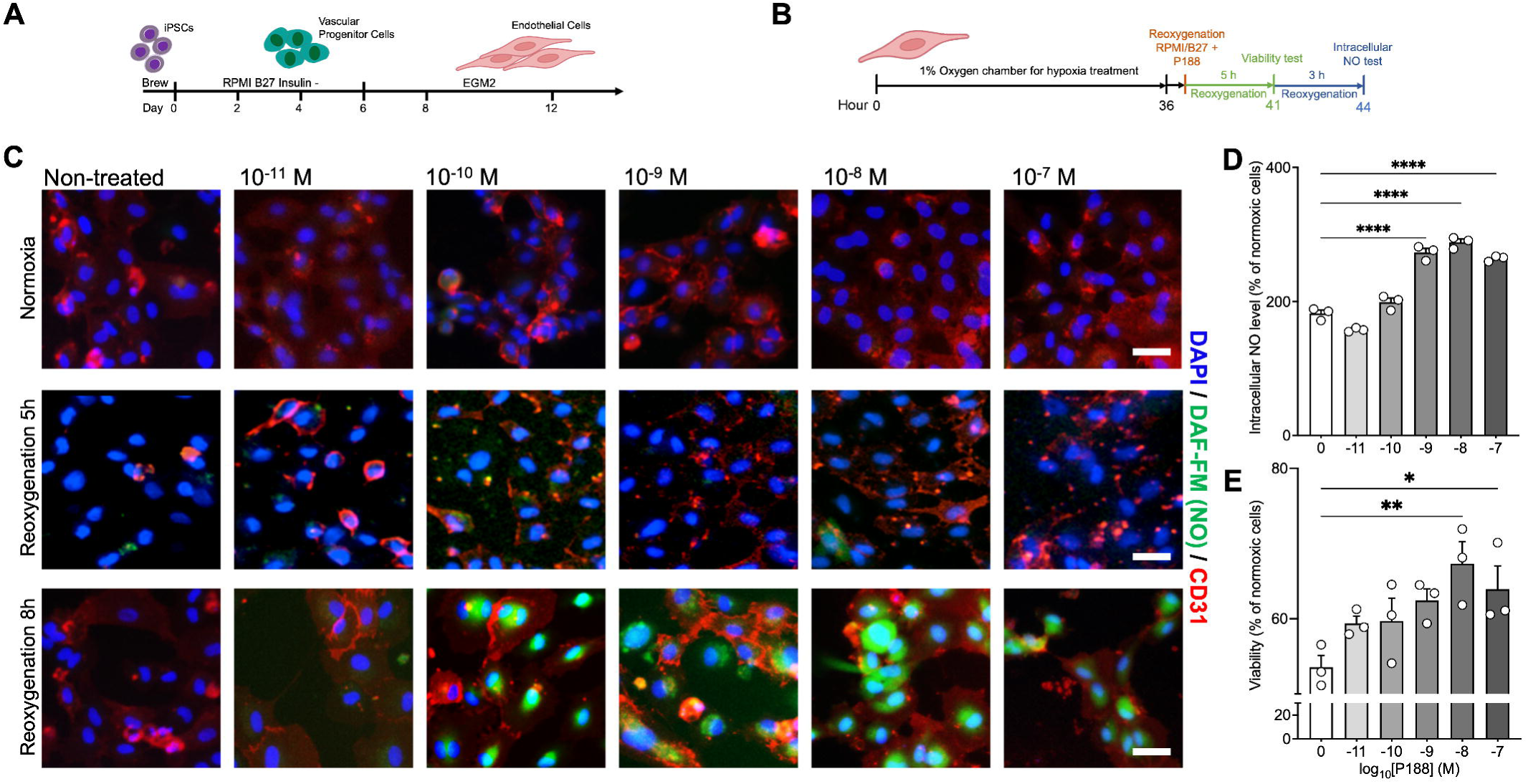
Effects of P188 on iPSC-ECs viability and intracellular NO production under simulated I/R conditions. (A) Schematic representation of the experimental timeline, detailing the differentiation protocol of iPSC-ECs followed by hypoxic exposure and subsequent reoxygenation. (B) Quantification of cell viability post-reperfusion in iPSC-ECs treated with or without P188. Cell viability values are normalized to normoxic control cells. (C) Quantification of intracellular NO production determined via DAF-FM staining. The values presented are normalized to normoxic control cells. (D) Representative images of intracellular NO staining, indicated by the DAF-FM dye. Positive intracellular NO production is depicted in green, nuclei are stained in blue, and the endothelial cell marker, CD31, is shown in red. The top row represents cells subjected to hypoxic treatment, while the bottom row shows corresponding cells under normal oxygen conditions, both treated with respective P188 concentrations; scale bars, 100 μm. Data are shown as mean ± SEM. Significance levels in comparison to the non-treated control group were determined using one-way ANOVA with Dunnett’s test: *P<0.05, **P<0.01, ****P<0.0001.

#### iPSC-CM apoptosis

To investigate the potential protective effect of P188 against programmed cell death, we quantified Annexin V-positive iPSC-derived CMs after 14 hours of reoxygenation. Flow cytometric analysis revealed that P188 reduced the proportion of apoptotic cells from 3.7% to 2.0% (***Fig. 2C***). The lowest level of Annexin V-positive portion was observed in the group treated with 10^-6^ M.

Together, these findings underscore the potential of P188 as a promising therapeutic agent for preserving iPSC-CM viability and function during hypoxia-reoxygenation events.

#### iPSC-EC function after hypoxia/reoxygenation experiments

Moreover, P188 treatment also exhibited a notable effect on the intracellular production of NO. Quantitative analyses revealed that the rate of NO production in ECs treated with P188 increased by approximately 50% relative to the control group that did not receive P188 treatment (***Fig. 3D***). The concentration-dependent effects of P188 on ECs were particularly pronounced at a concentration of 10 nM, where the most substantial improvements in both endothelial cell protection and intracellular NO production were observed after 8 hours of reoxygenation. Notably, the increase in NO production was not evident after 5 hours of reoxygenation but became significant after 8 hours ***(Fig. 3C)***. This temporal pattern suggests that the protective mechanisms activated by P188, leading to augmented NO production, may require a certain period before becoming fully operational within the cellular milieu. By enhancing EC survival and promoting NO release, P188 exhibits promising potential as a therapeutic agent to mitigate ischemic damage and improve cardiovascular outcomes.

#### iPSC-derived EC cell viability after hypoxia/reoxygenation experiments

Remarkably, P188 treatment led to a significant increase in EC viability. When compared to the control group without P188 treatment, the percentage of viable ECs increased from 55% to nearly 70% (***Fig. 3E***).

### P188 effect on myocardial I/R injury in rat isolated hearts

#### Left ventricular function and infarct size

At the end of the 120 minutes reperfusion phase, all ischemic hearts displayed a significant decrease in LVSP, LVDP, RPP, dP/dt_min_, dP/dt_max_ and CF and a significant increase in infarct size (***Fig. 4 A, 4C, 4E, 4G, and 4H***) compared to non-ischemic control experiments. P188 administered on reperfusion led to a significant decrease in LVEDP ***(Fig. 4B)***, increase in LVDP, RPP, dP/dt_min_ and decrease in infarct size ***(Fig. 4I)***. In P188 + L-NAME hearts, these improvements were abolished, while L-NAME itself had no significant effect by itself. There was no difference in HR among any of the groups ***(Fig. 4D)***.

**Fig. 4.**
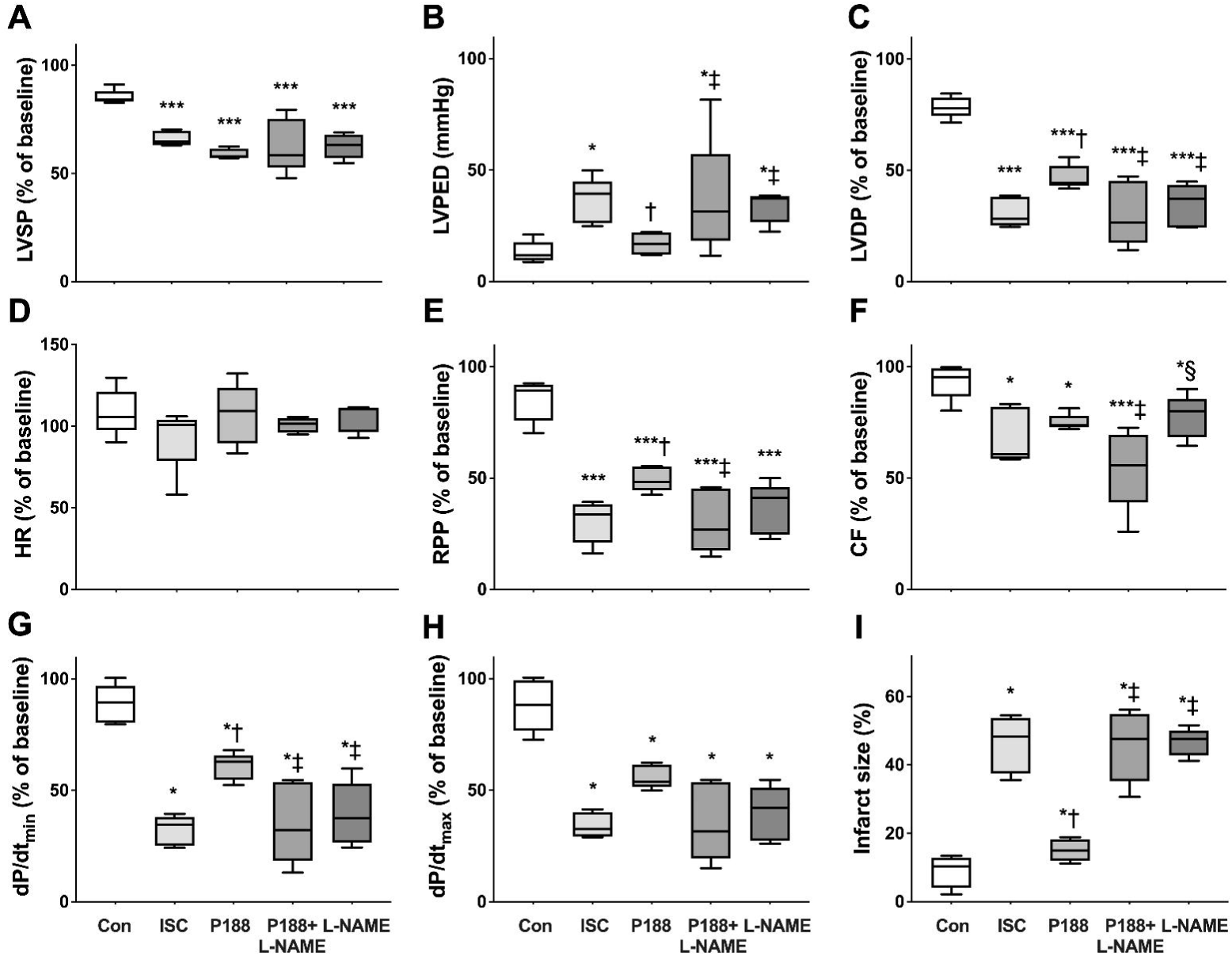
Effect of P188 on cardiac function and viability in rat isolated heart experiments. (A) Systolic (LVSP), (B) diastolic (LVEDP) and (C) developed left ventricular pressure (LVDP), (D) heart rate (HR), rate pressure product (RPP), (F) coronary flow (CF), (G) dP/dt_min_ and (H) dP/dt_max_, as indices of relaxation and contractility, respectively, and (I) infarct size are shown at 120 min of reperfusion. P188 significantly improved cardiac function and viability when compared to other groups undergoing ischemia/reperfusion. Notably, inhibition of nitric oxide synthase by Nω-Nitro-L-arginine methyl ester hydrochloride (L-NAME) abolished all effects of P188. Since not all data were normally distributed and had equal variance, we display them as box plots with median and interquartile range. Tests were considered statistically significant at P<0.05 (one symbol; P<0.01 two symbols; P<0.001 three symbols; P<0.0001 four symbols): *vs CON/control, † vs ISC, ‡ vs P188, § vs P188 + L-NAME.

#### NO measurements

P188 dose-dependently increased NO fluorescence in the absence of L-NAME. This inducible effect was completely abolished by L-NAME ***(Fig. 5A)***. In contrast, there was no increase in NO fluorescence produced in untreated control hearts (***Fig. 5B***).

**Fig. 5.**
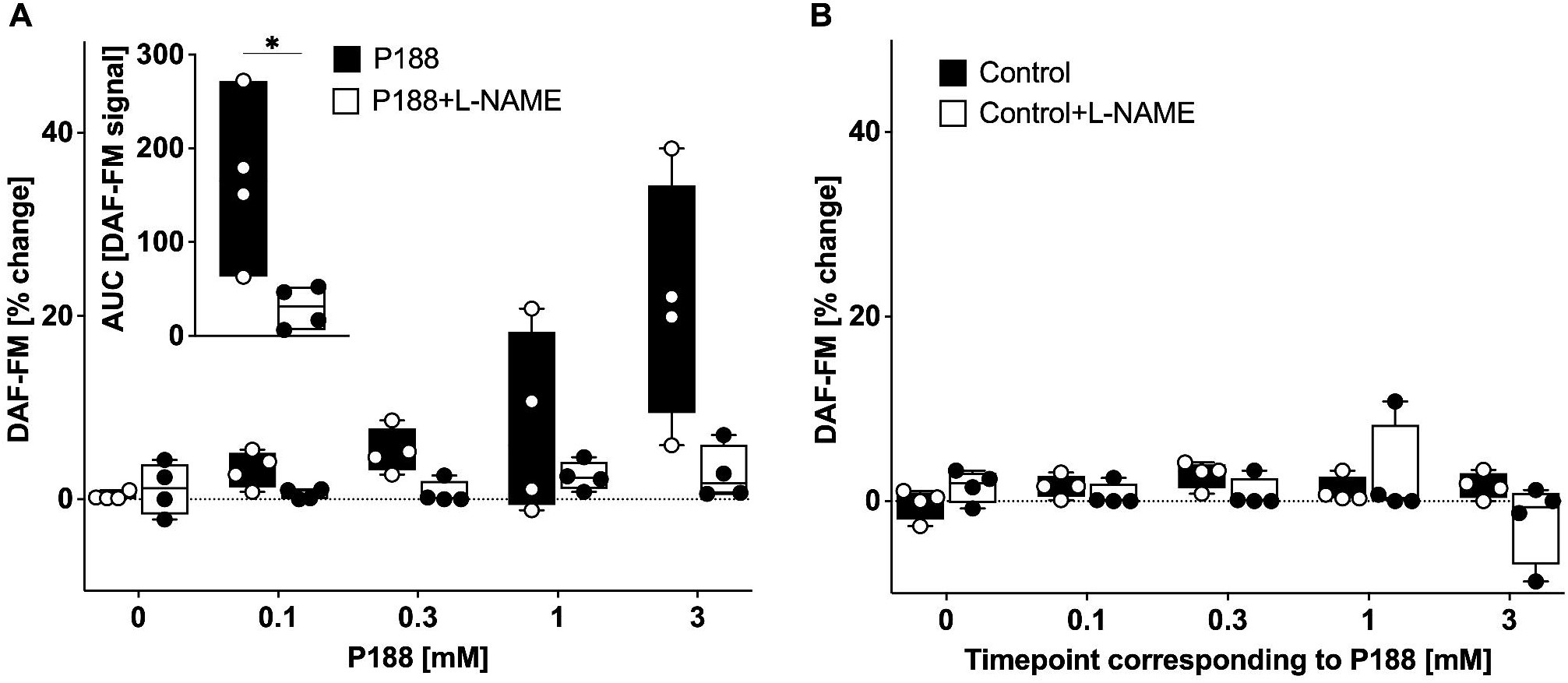
Nitric oxide measurement in rat isolated hearts at different P188 concentrations. (A) Fluorescence of the nitric oxide (NO)-sensitive dye 4-amino-5-methylamino-2′,7′-difluorofluorescein diacetate serving as an indicator of NO production revealed an increase after administration of P188 which was abolished by the nonspecific NO synthase inhibitor Nω-Nitro-L-arginine methyl ester hydrochloride (L-NAME). (A insert) In the absence of L-NAME, P188 increased the total amount of NO release, measured as the area under the curve (AUC) of the fluorescence signal; an effect which was abolished by L-NAME. (B) There was no increase in NO production in untreated control hearts. *P<0.05 vs P-188-NAME.

## Discussion

Our findings offer a novel insight into the multifaceted protective mechanisms of P188, highlighting its significant role beyond mere membrane stability. The ability of P188 to substantially increase NO production in ECs during reperfusion introduces a new dimension to its therapeutic potential. This increase in NO production, pivotal for vascular homeostasis and protection, suggests that P188’s benefits extend into modulating key signaling pathways involved in endothelial function and cardiovascular protection. P188-induced enhanced NO production may involve its amphipathic structure, which facilitates membrane stability and thereby potentially improves the functional status of membrane-bound enzymes such as eNOS. This hypothesis aligns with the observed delayed increase in NO production, suggesting a restoration of cellular functions because of preserved membrane integrity. These key results were validated in *ex-vivo* rat hearts, where P188 improved post-ischemia cardiac function in a NO-dependent manner. Moreover, P188 dose-dependently increased NO levels in non-ischemic beating hearts.

### Studies conducted in human induced pluripotent stem cell-derived CMs and ECs

We exposed both human iPSC-derived CMs and ECs to H/R stress and found that P188 dose-dependently protected both types of cells against reoxygenation injury. Human iPSC-derived CMs have been frequently used to study H/R injury [4, 16, 19, 21]. Our findings align with our previous studies showing that P188 can provide protection against reoxygenation injury [2] and that this protection depends on the dose of P188 [50]. These studies suggested that P188, similar to other poloxamers, may stabilize cell membranes or improve cell viability under stress conditions [26]. The mechanism is thought to involve the surfactant’s ability to integrate into the lipid bilayer of cell membranes, thereby reducing membrane stress and preventing leakage of intracellular components [17, 24]. Here, we utilized iPSC-CMs differentiated in regular RPMI/B27 medium, which made the cells relatively more resistant against hypoxic conditions compared to mature iPSC-CMs [47], requiring a longer hypoxia duration compared to our previous study [50]. Nevertheless, we could demonstrate that P188 given during reoxygenation resulted in a 25% reduction of H/R injury. We detected only a modest reduction in apoptosis by P188. This might in part be caused by the fact that apoptosis was measured late (i.e., 13 hours after reoxygenation) and emphasizes the acute effect of P188 on cell integrity upon reoxygenation.

Studies focusing on the examination of P188 in human tissue are still scarce, with only one prior publication conducted by Lin et al., which exclusively employed iPSC-CMs [28]. Remarkably, our investigation stands as the second study investigating the effects of P188 utilizing iPSCs, encompassing both iPSC-derived CMs and ECs. In contrast to the CM-only model employed by Lin et al., our bicellular model facilitates an enriched understanding of P188’s capabilities. Specifically, our research unveils the previously unexplored capacity of P188 to increase the secretion of NO in ECs, in addition to its recognized competence in membrane reparation evident by its effect on CM cell integrity [17, 22, 24].

### Langendorff experiments

#### Ischemia/reperfusion studies

Our findings unequivocally reveal a noteworthy amelioration in cardiac function across an extensive array of measured variables, accompanied by a considerable reduction in infarct size. The decrease observed in LVEDP among hearts subjected to P188 treatment primarily stems from enhanced relaxation during reperfusion. Similar investigations simulating myocardial infarction and congestive heart failure have consistently reported analogous outcomes, characterized by amplified LV end diastolic volume at reduced pressures, thereby yielding increased ejection fraction and cardiac function [2]. The state of constitutively overexpressing inducible NO synthase has previously demonstrated a comparable enhancement in LVEDP [64]. In our study, the observed improvement in LVEDP was entirely nullified by the administration of the non-specific NOS inhibitor L-NAME, indicating NO as key element in the protective mechanism exerted by P188. Furthermore, a strong cardioprotective effect has also been observed in an *in-vivo* porcine model when P188 was given immediately upon reperfusion following a prolonged coronary occlusion [23]. Given the statistical significance in I/R studies involving P188 and L-NAME, we are confident that P188, at least in part, is mediated by NO production.

#### NO studies

The results of the I/R studies prompted further investigation into the dose-dependent effects of P188 on NO production in naïve non-ischemic hearts. The initial studies elicited results supporting the hypothesis that P188 directly increases the rate and quantity of NO production in healthy, non-ischemic hearts. Additionally, preliminary studies (not shown) involving microscopic analysis of major heart vessels, revealed that hearts treated with P188 demonstrated significantly higher amounts of NO being produced [12]. Our results show an increased absolute amount of NO produced in hearts treated with increasing concentrations of P188 which was abolished in the presence of L-NAME (***Fig. 5A***). Moreover, the potential of P188 to act as an oxygen radical scavenger could also augment the availability of NO, as demonstrated by previous studies indicating that the mitigation of oxygen radical burden during reoxygenation via endothelial superoxide dismutase overexpression results in elevated NO availability [43]. These experiments provide further evidence the impact of P188 on NO signaling pathways.

It is known that during cardiac ischemia, the cellular membranes of cardiac ECs begin to degenerate following cellular swelling and extreme intracellular changes affecting multiple ion channels (for review see [20]). P188 has been shown to integrate into the damaged membranes to provide temporary protection and restore a physiological state until this lipid bilateral regenerates [24, 57]. Membrane integrity is imperative for prevention of microembolism of endothelial microaggregates leading to a no-reflow phenomenon [25]. Our findings suggest that P188 assists in sealing breaches in the degenerating membrane and thereby suppress release and aggregation of denaturated endothelial membrane proteins [41].

## Conclusion

Our approach of combining iPSCs with an *ex-vivo* animal model demonstrates that the amphiphilic copolymer P188 elicits cardioprotection against simulated I/R injury through coordinated membrane stabilization and an NO-dependent mechanisms. P188 protected human iPSC-derived CMs and ECs from H/R damage while boosting endothelial NO production. In rat isolated hearts, P188 improved post-ischemia cardiac function and viability in a NO-dependent manner and increased NO levels dose-dependently. These findings substantiate the previously reported potential of P188 – and of newer di-block derivatives [30] – as a novel therapy for I/R injury, warranting further evaluation and leveraging its dual capacity for membrane repair and NO potentiation during reperfusion [29, 31]. Cocultures with a close spatial relation between CMs and ECs will further help in elucidating the interplay of these two cell types in P188-induced cardioprotection [31].

### Author Contributions

Conceptualization GC, HFD, ZL, IYC, MLR and DO; methodology GC, HFD, IYC, MLR and DO; formal analysis CC, HFD, MLR and DO; data curation CC and HFD; writing—original draft preparation GC and HFD; writing—review and editing IYC, ZL, MLR, WJC, CB, DY; supervision MLR and DO; funding acquisition MLR, DY, DO. All authors have read and agreed to the published version of the manuscript.

### Funding

This research was funded – in part – by institutional funds, a Merit Review Award (I01 BX003482) from the U.S. Department of Veteran Affairs Biomedical Laboratory R&D Service to Dr. Riess, and NIH grant 5R01 HL123227 to Drs. Yannopoulos and Riess. Dr. Balzer received unrelated funding from the Deutsche Forschungsgemeinschaft (DFG, German Research Foundation) – Project BA 6287/1-1, Dr. Riess from a Transformative Project Award (962204) from the American Heart Association. Dr. Obal was funded through a TIP grant from the Maternal and Child Health Research Institute at Stanford University and a Startup Grant by the Department of Anesthesiology, Perioperative, and Pain Medicine.

### Institutional Review Board Statement

All human cell protocols for this study were approved by the Stanford University Human Subjects Research Institutional Review Board (IRB).

### Data Availability Statement

Any additional information required for reanalysis is available upon reasonable request and in strict accordance with funding guidelines.

## Acknowledgements

We thank Dr. Joseph C. Wu for providing access to SONY S18000 CM analysis tool. We thank both the Stanford Cardiovascular Institute (SCVI) Biobank and Stem Cell Core Facility of Genetics, Stanford University, which provided the human pluripotent cell lines SCVI-273 and SCVI-20. The authors would also like to thank Mr. James Heisner, Dr. Qunli Cheng, Dr. Matthew Olsen and Ms Michele Salzman for their technical assistance, and Amanda Chase for editing of the manuscript.

## Conflicts of Interest

The authors declare no conflict of interest.

---

**Supplement Table 1.** Reagents and Equipment.

| **Reagent / Equipment** | **Catalog Number / Model** | **Supplier** |
| --- | --- | --- |
| Annexin V-FITC/PI Apoptosis Detection Kit | A211 | Vazymebiotech |
| B27 Supplement (plus insulin) | A1486701 | Gibco |
| B27 Supplement (minus insulin) | A1895601 | Gibco |
| StemMACS™ iPS-Brew XF, human | 130-104-368 | Miltenyi Biotec |
| Calcein-AM | C3100MP | Invitrogen |
| CHIR99021, Free Base | C-6556 | LC Laboratories |
| DAF-FM diacetate | D23844 | Invitrogen |
| EGM™-2 Endothelial Cell Growth Medium-2 | CC-3162 | Lonza |
| Fibroblast Growth Factor-2 (rhFGF-2) | P09038 | PeproTech |
| Hoechst 33342 | H3570 | Invitrogen |
| KnockOut™ Serum Replacement | 10828010 | Gibco |
| Matrigel® Growth Factor Reduced Basement Matrix | 356231 | Corning |
| Nω-Nitro-L-arginine methyl ester hydrochloride | N5751 | Sigma Aldrich |
| P188 | K4894 | Sigma Aldrich |
| Propidium Iodide (PI) | P3566 | Invitrogen |
| ROCK Inhibitor Y-27632 | Y-5301 | LC Laboratories |
| RPMI 1640 Medium | 11875135 | Gibco |
| RPMI 1640 Medium (Glucose-free) | 11879020 | Gibco |
| Titer-Glo Luminescent Cell Viability Assay | G9241 | Promega |
| Trypsin-EDTA (0.25%) | 25200056 | ThermoFisher Scientific |
| 2,3,5-triphenyltetrazolium chloride | T8877 | Sigma Aldrich |
| UltraPure 0.5M EDTA | 15575-038 | Invitrogen |
| Wnt-C59 (C59) | S7037 | Selleck Chemical |
| **Equipment** |  |  |
| AC converter NI USB-6343 |  | National Instruments Corporation |
| Hypoxia Incubator Chamber | 27310 | STEMCELL Technologies |
| Cell Motion Imaging System | SI8000 | Sony Biotechnology |
| Microscope | ECHO-Revolve | ECHO |
| Flow Cytometer | Accuri C6 | BD Biosciences |
| Multi Plate Reader | GloMax | Promega |
| Oxygen Detector 0-30% | FD-90A-O2 | Forensics Detectors |
| Ultrasonic Flowmeter | T106X | Transonic Systems |
| **Software** |  |  |
| Image J | Ver. 2.14.0 | NIH |
| GraphPad Prism | Ver. 10 | Graphpad Software, LLC. |
| Labview | Ver. 2014 | National Instruments |
| R studio | Ver. 2023.06.2 | Posit Software |
| SI8000 | Ver. 1.03.0008 | Sony Biotechnology |

## Notes

### Competing Interest Statement

The authors have declared no competing interest.

## Literature

1. Bao H-J, Wang T, Zhang M-Y, Liu R, Dai D-K, Wang Y-Q, Wang L, Zhang L, Gao Y-Z, Qin Z-H, Chen X-P, Tao L-Y (2012) Poloxamer-188 attenuates TBI-induced blood-brain barrier damage leading to decreased brain edema and reduced cellular death. Neurochem Res 37:2856–2867. doi: 10.1007/s11064-012-0880-4

2. Bartos JA, Matsuura TR, Tsangaris A, Olson M, McKnite SH, Rees JN, Haman K, Shekar KC, Riess ML, Bates FS, Metzger JM, Yannopoulos D (2016) Intracoronary Poloxamer 188 Prevents Reperfusion Injury in a Porcine Model of ST-Segment Elevation Myocardial Infarction. JACC Basic Transl Sci 1:224–234. doi: 10.1016/j.jacbts.2016.04.001

3. Bates FS, Hillmyer MA, Lodge TP, Bates CM, Delaney KT, Fredrickson GH (2012) Multiblock polymers: panacea or Pandora’s box? Science 336:434–440. doi: 10.1126/science.1215368

4. Brodarac A, Šarić T, Oberwallner B, Mahmoodzadeh S, Neef K, Albrecht J, Burkert K, Oliverio M, Nguemo F, Choi Y-H, Neiss WF, Morano I, Hescheler J, Stamm C (2015) Susceptibility of murine induced pluripotent stem cell-derived cardiomyocytes to hypoxia and nutrient deprivation. Stem Cell Res Ther 6:83. doi: 10.1186/s13287-015-0057-6

5. Burridge PW, Matsa E, Shukla P, Lin ZC, Churko JM, Ebert AD, Lan F, Diecke S, Huber B, Mordwinkin NM, Plews JR, Abilez OJ, Cui B, Gold JD, Wu JC (2014) Chemically defined generation of human cardiomyocytes. Nat Methods 11:855–860. doi: 10.1038/nmeth.2999

6. Chatterjee S, Fisher AB (2014) Mechanotransduction in the endothelium: role of membrane proteins and reactive oxygen species in sensing, transduction, and transmission of the signal with altered blood flow. Antioxid Redox Signal 20:899–913. doi: 10.1089/ars.2013.5624

7. Chen H-P, Liao Z-P, Huang Q-R, He M (2009) Sodium ferulate attenuates anoxia/reoxygenation-induced calcium overload in neonatal rat cardiomyocytes by NO/cGMP/PKG pathway. Eur J Pharmacol 603:86–92. doi: 10.1016/j.ejphar.2008.12.003

8. Chiu J-J, Chien S (2011) Effects of disturbed flow on vascular endothelium: pathophysiological basis and clinical perspectives. Physiol Rev 91:327–387. doi: 10.1152/physrev.00047.2009

9. Coute RA, Nathanson BH, Mader TJ, McNally B, Kurz MC (2021) Trend analysis of disability-adjusted life years following adult out-of-hospital cardiac arrest in the United States: A study from the CARES Surveillance Group. Resuscitation 163:124–129. doi: 10.1016/j.resuscitation.2020.10.048

10. Davis ME, Cai H, McCann L, Fukai T, Harrison DG (2003) Role of c-Src in regulation of endothelial nitric oxide synthase expression during exercise training. Am J Physiol Heart Circ Physiol 284:H1449–53. doi: 10.1152/ajpheart.00918.2002

11. Dimmeler S, Fleming I, Fisslthaler B, Hermann C, Busse R, Zeiher AM (1999) Activation of nitric oxide synthase in endothelial cells by Akt-dependent phosphorylation. Nature 399:601–605. doi: 10.1038/21224

12. Douglas HF, Salzman MM, Hackel BJ, Bartos JA, Yannopolous D, Riess ML (2018) Dose Dependent Nitric Oxide Production by Poloxamer 188 in Rat Isolated Hearts. The FASEB Journal

13. Dynnik VV, Grishina EV, Fedotcheva NI (2020) The mitochondrial NO-synthase/guanylate cyclase/protein kinase G signaling system underpins the dual effects of nitric oxide on mitochondrial respiration and opening of the permeability transition pore. FEBS J 287:1525–1536. doi: 10.1111/febs.15090

14. Erkens R, Suvorava T, Kramer CM, Diederich LD, Kelm M, Cortese-Krott MM (2017) Modulation of local and systemic heterocellular communication by mechanical forces: A role of endothelial nitric oxide synthase. Antioxid Redox Signal 26:917–935. doi: 10.1089/ars.2016.6904

15. Fleming I, Fisslthaler B, Dimmeler S, Kemp BE, Busse R (2001) Phosphorylation of Thr(495) regulates Ca(2+)/calmodulin-dependent endothelial nitric oxide synthase activity. Circ Res 88:E68–75. doi: 10.1161/hh1101.092677

16. Funcke S, Werner TR, Hein M, Ulmer BM, Hansen A, Eschenhagen T, Hirt MN (2020) Effects of the Delta Opioid Receptor Agonist DADLE in a Novel Hypoxia-Reoxygenation Model on Human and Rat-Engineered Heart Tissue: A Pilot Study. Biomolecules. doi: 10.3390/biom10091309

17. Goliaei A, Lau EY, Adhikari U, Schwegler E, Berkowitz ML (2016) Behavior of P85 and P188 Poloxamer Molecules: Computer Simulations Using United-Atom Force-Field. J Phys Chem B 120:8631–8641. doi: 10.1021/acs.jpcb.6b03030

18. Gu J-H, Ge J-B, Li M, Xu H-D, Wu F, Qin Z-H (2013) Poloxamer 188 protects neurons against ischemia/reperfusion injury through preserving integrity of cell membranes and blood brain barrier. PLoS ONE 8:e61641. doi: 10.1371/journal.pone.0061641

19. Häkli M, Kreutzer J, Mäki A-J, Välimäki H, Lappi H, Huhtala H, Kallio P, Aalto-Setälä K, Pekkanen-Mattila M (2021) Human induced pluripotent stem cell-based platform for modeling cardiac ischemia. Sci Rep 11:4153. doi: 10.1038/s41598-021-83740-w

20. Heusch G (2016) The coronary circulation as a target of cardioprotection. Circ Res 118:1643–1658. doi: 10.1161/CIRCRESAHA.116.308640

21. Hidalgo A, Glass N, Ovchinnikov D, Yang S-K, Zhang X, Mazzone S, Chen C, Wolvetang E, Cooper-White J (2018) Modelling ischemia-reperfusion injury (IRI) in vitro using metabolically matured induced pluripotent stem cell-derived cardiomyocytes. APL Bioengineering 2:026102. doi: 10.1063/1.5000746

22. Houang EM, Bartos J, Hackel BJ, Lodge TP, Yannopoulos D, Bates FS, Metzger JM (2019) Cardiac muscle membrane stabilization in myocardial reperfusion injury. JACC Basic Transl Sci 4:275–287. doi: 10.1016/j.jacbts.2019.01.009

23. Juneman EB, Saleh L, Lancaster JJ, Thai HM, Markham B, Goldman S (2012) The effects of poloxamer-188 on left ventricular function in chronic heart failure after myocardial infarction. J Cardiovasc Pharmacol 60:293–298. doi: 10.1097/FJC.0b013e31825f6f88

24. Kim M, Heinrich F, Haugstad G, Yu G, Yuan G, Satija SK, Zhang W, Seo HS, Metzger JM, Azarin SM, Lodge TP, Hackel BJ, Bates FS (2020) Spatial Distribution of PEO-PPO-PEO Block Copolymer and PEO Homopolymer in Lipid Bilayers. Langmuir 36:3393–3403. doi: 10.1021/acs.langmuir.9b03208

25. Kloner RA, Ganote CE, Jennings RB (1974) The “no-reflow” phenomenon after temporary coronary occlusion in the dog. J Clin Invest 54:1496–1508. doi: 10.1172/JCI107898

26. Kwiatkowski TA, Rose AL, Jung R, Capati A, Hallak D, Yan R, Weisleder N (2020) Multiple poloxamers increase plasma membrane repair capacity in muscle and nonmuscle cells. Am J Physiol, Cell Physiol 318:C253–C262. doi: 10.1152/ajpcell.00321.2019

27. Lian X, Hsiao C, Wilson G, Zhu K, Hazeltine LB, Azarin SM, Raval KK, Zhang J, Kamp TJ, Palecek SP (2012) Robust cardiomyocyte differentiation from human pluripotent stem cells via temporal modulation of canonical Wnt signaling. Proc Natl Acad Sci USA 109:E1848–57. doi: 10.1073/pnas.1200250109

28. Lin B, Li Y, Han L, Kaplan AD, Ao Y, Kalra S, Bett GCL, Rasmusson RL, Denning C, Yang L (2015) Modeling and study of the mechanism of dilated cardiomyopathy using induced pluripotent stem cells derived from individuals with Duchenne muscular dystrophy. Dis Model Mech 8:457–466. doi: 10.1242/dmm.019505

29. Li Z, Barajas M, Oyama T, Riess ML (2023) Abstract 363: endothelial nitric oxide mediates protection of isolated cardiomyocytes by poloxamer 188 against hypoxia reoxygenation injury. Circulation. doi: 10.1161/circ.148.suppl_1.363

30. Li Z, Gupta MK, Barajas MB, Oyama T, Duvall CL, Riess ML (2023) Newly Developed Di-Block Copolymer-Based Cell Membrane Stabilizers Protect Mouse Coronary Artery Endothelial Cells against Hypoxia/Reoxygenation Injury. Cells. doi: 10.3390/cells12101394

31. Li Z, Hampton MJW, Barajas MB, Riess ML (2021) Development of a Cell Co-Culture Model to Mimic Cardiac Ischemia/Reperfusion In Vitro. J Vis Exp. doi: 10.3791/62913

32. Luo C-L, Chen X-P, Li L-L, Li Q-Q, Li B-X, Xue A-M, Xu H-F, Dai D-K, Shen Y-W, Tao L-Y, Zhao Z-Q (2013) Poloxamer 188 attenuates in vitro traumatic brain injury-induced mitochondrial and lysosomal membrane permeabilization damage in cultured primary neurons. J Neurotrauma 30:597–607. doi: 10.1089/neu.2012.2425

33. Luo J, Obal D, Dimova N, Tang X-L, Rokosh G (2013) Cardiac myocyte-specific transgenic ecSOD targets mitochondria to protect against Ca(2+) induced permeability transition. Front Physiol 4:295. doi: 10.3389/fphys.2013.00295

34. Madathil RJ, Hira RS, Stoeckl M, Sterz F, Elrod JB, Nichol G (2016) Ischemia reperfusion injury as a modifiable therapeutic target for cardioprotection or neuroprotection in patients undergoing cardiopulmonary resuscitation. Resuscitation 105:85–91. doi: 10.1016/j.resuscitation.2016.04.009

35. Martindale JJ, Metzger JM (2014) Uncoupling of increased cellular oxidative stress and myocardial ischemia reperfusion injury by directed sarcolemma stabilization. J Mol Cell Cardiol 67:26–37. doi: 10.1016/j.yjmcc.2013.12.008

36. Maskarinec SA, Hannig J, Lee RC, Lee KYC (2002) Direct observation of poloxamer 188 insertion into lipid monolayers. Biophys J 82:1453–1459. doi: 10.1016/S0006-3495(02)75499-4

37. Moloughney JG, Weisleder N (2012) Poloxamer 188 (p188) as a membrane resealing reagent in biomedical applications. Recent Pat Biotechnol 6:200–211. doi: 10.2174/1872208311206030200

38. Moncada S, Higgs EA (2006) Nitric oxide and the vascular endothelium. Handb Exp Pharmacol 213–254. doi: 10.1007/3-540-32967-6_7

39. Mosqueira M, Konietzny R, Andresen C, Wang C, H A Fink R (2021) Cardiomyocyte depolarization triggers NOS-dependent NO transient after calcium release, reducing the subsequent calcium transient. Basic Res Cardiol 116:18. doi: 10.1007/s00395-021-00860-0

40. Murphy AD, McCormack MC, Bichara DA, Nguyen JT, Randolph MA, Watkins MT, Lee RC, Austen WG (2010) Poloxamer 188 protects against ischemia-reperfusion injury in a murine hind-limb model. Plast Reconstr Surg 125:1651–1660. doi: 10.1097/PRS.0b013e3181ccdbef

41. Mustafi D, Smith CM, Makinen MW, Lee RC (2008) Multi-block poloxamer surfactants suppress aggregation of denatured proteins. Biochim Biophys Acta 1780:7–15. doi: 10.1016/j.bbagen.2007.08.017

42. Nichol G, Soar J (2010) Regional cardiac resuscitation systems of care. Curr Opin Crit Care 16:223–230. doi: 10.1097/MCC.0b013e32833985b5

43. Obal D, Dai S, Keith R, Dimova N, Kingery J, Zheng Y-T, Zweier J, Velayutham M, Prabhu SD, Li Q, Conklin D, Yang D, Bhatnagar A, Bolli R, Rokosh G (2012) Cardiomyocyte-restricted overexpression of extracellular superoxide dismutase increases nitric oxide bioavailability and reduces infarct size after ischemia/reperfusion. Basic Res Cardiol 107:305. doi: 10.1007/s00395-012-0305-1

44. Osawa M, Masuda M, Kusano K, Fujiwara K (2002) Evidence for a role of platelet endothelial cell adhesion molecule-1 in endothelial cell mechanosignal transduction: is it a mechanoresponsive molecule? J Cell Biol 158:773–785. doi: 10.1083/jcb.200205049

45. Paik DT, Tian L, Lee J, Sayed N, Chen IY, Rhee S, Rhee J-W, Kim Y, Wirka RC, Buikema JW, Wu SM, Red-Horse K, Quertermous T, Wu JC (2018) Large-Scale Single-Cell RNA-Seq Reveals Molecular Signatures of Heterogeneous Populations of Human Induced Pluripotent Stem Cell-Derived Endothelial Cells. Circ Res 123:443–450. doi: 10.1161/CIRCRESAHA.118.312913

46. Patel VH, Brack KE, Coote JH, Ng GA (2008) A novel method of measuring nitric-oxide-dependent fluorescence using 4,5-diaminofluorescein (DAF-2) in the isolated Langendorff-perfused rabbit heart. Pflugers Arch 456:635–645. doi: 10.1007/s00424-007-0440-y

47. 47. Peters MC, Maas RGC, van Adrichem I, Doevendans PAM, Mercola M, Šarić T, Buikema JW, van Mil A, Chamuleau SAJ, Sluijter JPG, Hnatiuk AP, Neef K (2022) Metabolic Maturation Increases Susceptibility to Hypoxia-induced Damage in Human iPSC-derived Cardiomyocytes. Stem Cells Transl Med 11:1040–1051. doi: 10.1093/stcltm/szac061

48. Riess ML, Rhodes SS, Stowe DF, Aldakkak M, Camara AKS (2009) Comparison of cumulative planimetry versus manual dissection to assess experimental infarct size in isolated hearts. J Pharmacol Toxicol Methods 60:275–280. doi: 10.1016/j.vascn.2009.05.012

49. Roux E, Bougaran P, Dufourcq P, Couffinhal T (2020) Fluid shear stress sensing by the endothelial layer. Front Physiol 11:861. doi: 10.3389/fphys.2020.00861

50. Salzman MM, Bartos JA, Yannopoulos D, Riess ML (2020) Poloxamer 188 Protects Isolated Adult Mouse Cardiomyocytes from Reoxygenation Injury. Pharmacol Res Perspect 8:e00639. doi: 10.1002/prp2.639

51. Schulz R, Kelm M, Heusch G (2004) Nitric oxide in myocardial ischemia/reperfusion injury. Cardiovasc Res 61:402–413. doi: 10.1016/j.cardiores.2003.09.019

52. Serbest G, Horwitz J, Jost M, Barbee K (2006) Mechanisms of cell death and neuroprotection by poloxamer 188 after mechanical trauma. FASEB J 20:308–310. doi: 10.1096/fj.05-4024fje

53. Shidham S, Nabbi R, Camara AKS, Riess ML (2011) Development of Automated Infarct Size Measurement in TTC Stained Rat Isolated Hearts after Global Ischemia/Reperfusion. FASEB J. doi: 10.1096/fasebj.25.1_supplement.1130.2

54. Shyy JY-J, Chien S (2002) Role of integrins in endothelial mechanosensing of shear stress. Circ Res 91:769–775. doi: 10.1161/01.res.0000038487.19924.18

55. Strimaityte D, Tu C, Yanez A, Itzhaki I, Wu H, Wu JC, Yang H (2022) Contractility and Calcium Transient Maturation in the Human iPSC-Derived Cardiac Microfibers. ACS Appl Mater Interfaces 14:35376–35388. doi: 10.1021/acsami.2c07326

56. Stumpe T, Decking UK, Schrader J (2001) Nitric oxide reduces energy supply by direct action on the respiratory chain in isolated cardiomyocytes. Am J Physiol Heart Circ Physiol 280:H2350–6. doi: 10.1152/ajpheart.2001.280.5.H2350

57. Torres Filho IP, Torres LN, Salgado C, Dubick MA (2017) Novel adjunct drugs reverse endothelial glycocalyx damage after hemorrhagic shock in rats. Shock 48:583–589. doi: 10.1097/SHK.0000000000000895

58. Townsend D, Turner I, Yasuda S, Martindale J, Davis J, Shillingford M, Kornegay JN, Metzger JM (2010) Chronic administration of membrane sealant prevents severe cardiac injury and ventricular dilatation in dystrophic dogs. J Clin Invest 120:1140–1150. doi: 10.1172/JCI41329

59. Tzima E, Irani-Tehrani M, Kiosses WB, Dejana E, Schultz DA, Engelhardt B, Cao G, DeLisser H, Schwartz MA (2005) A mechanosensory complex that mediates the endothelial cell response to fluid shear stress. Nature 437:426–431. doi: 10.1038/nature03952

60. Walters TJ, Mase VJ, Roe JL, Dubick MA, Christy RJ (2011) Poloxamer-188 reduces muscular edema after tourniquet-induced ischemia-reperfusion injury in rats. J Trauma 70:1192–1197. doi: 10.1097/TA.0b013e318217879a

61. Wang T, Chen X, Wang Z, Zhang M, Meng H, Gao Y, Luo B, Tao L, Chen Y (2015) Poloxamer-188 can attenuate blood-brain barrier damage to exert neuroprotective effect in mice intracerebral hemorrhage model. J Mol Neurosci 55:240–250. doi: 10.1007/s12031-014-0313-8

62. Wang Z, Yang N, Hou Y, Li Y, Yin C, Yang E, Cao H, Hu G, Xue J, Yang J, Liao Z, Wang W, Sun D, Fan C, Zheng L (2023) L-Arginine-Loaded Gold Nanocages Ameliorate Myocardial Ischemia/Reperfusion Injury by Promoting Nitric Oxide Production and Maintaining Mitochondrial Function. Adv Sci (Weinh) e2302123. doi: 10.1002/advs.202302123

63. Wei T-T, Chandy M, Nishiga M, Zhang A, Kumar KK, Thomas D, Manhas A, Rhee S, Justesen JM, Chen IY, Wo H-T, Khanamiri S, Yang JY, Seidl FJ, Burns NZ, Liu C, Sayed N, Shie J-J, Yeh C-F, Yang K-C, Lau E, Lynch KL, Rivas M, Kobilka BK, Wu JC (2022) Cannabinoid receptor 1 antagonist genistein attenuates marijuana-induced vascular inflammation. Cell 185:2387–2389. doi: 10.1016/j.cell.2022.06.006

64. West MB, Rokosh G, Obal D, Velayutham M, Xuan Y-T, Hill BG, Keith RJ, Schrader J, Guo Y, Conklin DJ, Prabhu SD, Zweier JL, Bolli R, Bhatnagar A (2008) Cardiac myocyte-specific expression of inducible nitric oxide synthase protects against ischemia/reperfusion injury by preventing mitochondrial permeability transition. Circulation 118:1970–1978. doi: 10.1161/CIRCULATIONAHA.108.791533

65. Williams JG, Ojaimi C, Qanud K, Zhang S, Xu X, Recchia FA, Hintze TH (2008) Coronary nitric oxide production controls cardiac substrate metabolism during pregnancy in the dog. Am J Physiol Heart Circ Physiol 294:H2516–23. doi: 10.1152/ajpheart.01196.2007

66. Yasuda S, Townsend D, Michele DE, Favre EG, Day SM, Metzger JM (2005) Dystrophic heart failure blocked by membrane sealant poloxamer. Nature 436:1025–1029. doi: 10.1038/nature03844

